# *E. coli* cells advance into phase-separated (biofilm-simulating) extracellular polymeric substance containing DNA, HU, and lipopolysaccharide

**DOI:** 10.1101/2024.06.18.599655

**Authors:** Archit Gupta, Purnananda Guptasarma

**Affiliations:** Centre for Protein Science, Design and Engineering, Indian Institute of Science Education and Research (IISER) Mohali, Knowledge City, Sector 81, SAS Nagar, Punjab 140306, India; Department of Biological Sciences; Indian Institute of Science Education and Research (IISER) Mohali, Knowledge City, Sector 81, SAS Nagar, Punjab 140306, India

## Abstract

We have earlier shown that HU, a nucleoid-associated protein, uses its DNA-binding surfaces to bind to bacterial outer-membrane lipopolysachharide (LPS), with this explaining how HU act as a potential glue for the adherence of bacteria to DNA, e.g., in biofilms. We have also earlier shown that HU and DNA together condense into a state of liquid-liquid phase separation (LLPS) both within, and outside, bacterial cells. Here, we report that HU and free LPS also condense into a state of phase separation, with coacervates of HU, DNA and free LPS being less liquid-like than condensates of HU and DNA alone. *E. coli* cells bearing surface LPS and also shedding LPS, are shown to adhere to (as well as enter into) condensates of HU and DNA. HU thus appears to play a role in maintaining both an intracellular state of phase separation involving genomic nucleoids that are phase-separated from the cytoplasm, and an extracellular state of phase separation involving coacervates of extracellular DNA, HU and LPS, in the extracellular polymeric substances (EPS) of biofilms, in which LPS content is shown to modulate liquidity.

**Importance:** Understanding biofilm genesis and nature are crucial to understanding how to deal with the bacterial resistance to antibiotics that develops eventually in persistent biofilms. This study, together with two other recent landmark studies from our group, elucidates a novel aspect of the extracellular polymeric substance (EPS) of an *Escherichia coli* biofilm, by creating a simulacrum of the EPS and demonstrating that its formation involves liquid-liquid phase separation (LLPS) by its component HU, DNA, and lipopolysaccharide (LPS), with LPS determining the liquidity of the EPS simulacrum. The findings provide insight into the physical nature of biofilms but also suggests that the interplay of HU, DNA, and LPS facilitates the structural integrity and functional dynamics of biofilms. These findings are a stepping stone to the eventual development of strategies to disrupt biofilm.

## Introduction

*Escherichia coli* forms conglomerations of cells called biofilms, within which cells are held together by an extracellular polymeric substance (EPS) that is rich in nucleic acids [1]. The nucleic acids include both RNA and extracellular DNA [2] that is bound to DNA-binding proteins [3]. One abundant protein in the EPS is a non-sequence-specific, nucleoid-associated, DNA-binding protein known as HU [4]. The EPS (and consequently, biofilms too) has been shown to undergo disintegration through the action of nucleases that degrade the extracellular DNA matrix, as well as through the action of upon antibodies that are cognate to the DNA-binding tips of DNABII proteins such as HU [5–7]. This suggests that the extracellular DNA somehow requires DNABII proteins like HU to be present upon it, for it to hold *E. coli* cells together in the biofilm. Some years ago, we demonstrated that HU uses its canonical and non-canonical DNA-binding surfaces to bind to lipopolysaccharide (LPS) that is present upon the outer membranes of *E*. coli, with HU appearing to recognize the hexose sugar-linked phosphate groups on the headgroup of LPS as a proxy for the pentose sugar-linked phosphate groups on the backbone of DNA [8]. The demonstration that HU is a charge-neutralizing glue which allows the negatively-charged surface of a bacterium to bind to the negatively-charged surface of DNA suggested mechanisms for both (i) the embedment of bacteria within a matrix of DNA, and (ii) the generation of extracellular DNA to sustain the growth of the biofilm, through the occasional lysis of cells that tug at each other. Notably, the explosive lysis of bacterial cells that appear to tug at each other has been observed experimentally, and also shown to generate extracellular DNA [9,10]. Therefore, our identification of HU as a molecular glue answered questions both about how negatively-charged *E. coli* cells coexist with negatively charged DNA [8,11] in biofilms, and also about how extracellular DNA comes to exist in biofilms.

Independently, numerous experimental and computational (simulation-based) studies have suggested over the years that biofilms could be the result of phase transitions involving either liquid-liquid phase separation (LLPS), or the formation of a gel-like substance [12–14]. Theoretically-speaking, phase transitions involving the generation of varying degrees of liquid-like, gel-like, or solid-like behaviour, could most certainly hold benefits for the formation, maintenance and differentiation of biofilms, since scope for differences between different regions of a biofilm (e.g., inner and outer regions, or older and newer regions) could help a biofilm to become a complex collection of living cells, dead cells and EPS derived from dead cells. Multiple hypotheses abound regarding how biofilms form through the phase-separation of bacterial populations from their immediate environments [14]. Given that a biofilm’s EPS is rich in DNA and DNA-binding proteins like HU, it is significant that we have recently demonstrated that HU and DNA crowd each other into a liquid-liquid phase-separated (LLPS) state, both within bacterial genomic nucleoids, and *in vitro* [15]. Thus, it is plausible that biofilms are sustained through the cell lysis-based disgorgement of pre-formed biomolecular (HU-DNA) condensates derived from genomic nucleoids.

In other words, our previous work showing (i) that HU binds to LPS [ref], and (ii) that HU and DNA crowd each other into LLPS states [15], seems to suggest that LLPS condensates of HU and DNA constitute the matrix of a biofilm, and that bacterial cells are embedded in this matrix, through the binding of bacterial cell-surface LPS to the HU that is present within extracellular LLPS condensates. This makes it necessary to explicitly examine whether *E. coli* cells show a preference for existing within condensates of HU and DNA, and also examine the role that is played in such condensates by the free LPS that is naturally shed by bacterial cells. Here, we demonstrate the following: (i) that HU also phase-separates with free LPS, as it does with DNA, but that HU-LPS condensates are less liquid-like than HU-DNA condensates; (ii) that HU, DNA and free LPS coacervate to form mixed condensates that are less liquid-like that HU-DNA condensates; and (iii) that bacteria readily advance into HU-DNA condensates.

## Results

### LPS and HU associate to undergo biomolecular condensation

We have previously shown that HU undergoes phase separation with DNA under physiological conditions (13, 14), using physiological concentrations of HU and DNA base pairs. Below, we demonstrate HU’s ability to undergo phase separation with LPS, under similar conditions, with one notable exception. With DNA, we had shown that HU forms condensates even in the complete absence of any macromolecular crowding agent such as polyethylene glycol. However, to reduce the amount of synthetic DNA used, we did use a nominal concentration of 2 % PEG in all experiments carried out thereafter with HU and DNA. Here, with LPS, we found that the mutual crowding of HU-B and LPS is not as effective, presumably owing to the fact that each LPS molecule contains only two sugar-phosphate moieties for the binding of HU-B, unlike DNA which contains many more sugar-phosphate moieties for HU-B binding. Thus, there was no condensation in the absence of PEG. In the presence of a nominal PEG concentration of 4 %, however, addition of LPS to HU, led to the formation of spherical biomolecular droplet-like condensates.

Figure 1A shows three tubes containing 100 μM HU-B in 150 mM salt (KCl), 20 mM Tris, and 4 % PEG 6000, at pH 7.4, (i) in the complete absence of DNA or LPS, (ii) in the presence of LPS alone, and (ii) in the presence of cruciform 4-way junction (4WJ) DNA alone. For these experiments, similar w/v concentrations of LPS and DNA were used, because the average molecular weight of LPS, which is 50-100 kDa [17–19], is similar to the molecular weight of 4WJ DNA [15], which is ∼46.5 kDa. Cruciform DNA was used in these experiments both because HU-B displays a high affinity for cruciforms [20], which helps to reduce the amounts of synthetic DNA used, and also because extracellular DNA has been reported to be rich in cruciform DNA [21]. The tube containing HU-B alone displays no turbidity. Tubes containing either HU-B and DNA, or HU-B and LPS, show turbidity. The degree of turbidity in the tube containing HU and LPS is significantly lower than that seen in the tube containing HU-B and DNA. The LLPS nature of the entities giving rise to the turbidity was verified through fluorescence confocal microcopy. Figure 1B shows control confocal microscopic images of HU-B alone (100 μM), or LPS alone (0.4 mg/ml), revealing that no phase separation is seen in either instance, using the aforementioned buffer conditions. Figure 1C reveals that mixtures of HU-B and LPS form spherical condensates displaying both LPS-derived (red) and HU-derived (green) fluorescence. The Mander’s and Pearson’s coefficients of colocalization were calculated for HU-B and LPS (Supplementary Figure S1), and the colocalization was found to be significant, using all parameters used. The spherical morphology of the HU-LPS condensates was further validated using DIC imaging (Supplementary Figure S2). The presence of LPS inside the spherical condensates establishes that LPS acts not simply as a crowding agent that fails to enter condensates, but rather as an entity that actively engages with HU-B to undergo complex coacervation. This is clearly established by the overlap in the fluorescence signals derived from HU-B and LPS, in the spherical condensates formed.

**Figure 1.**
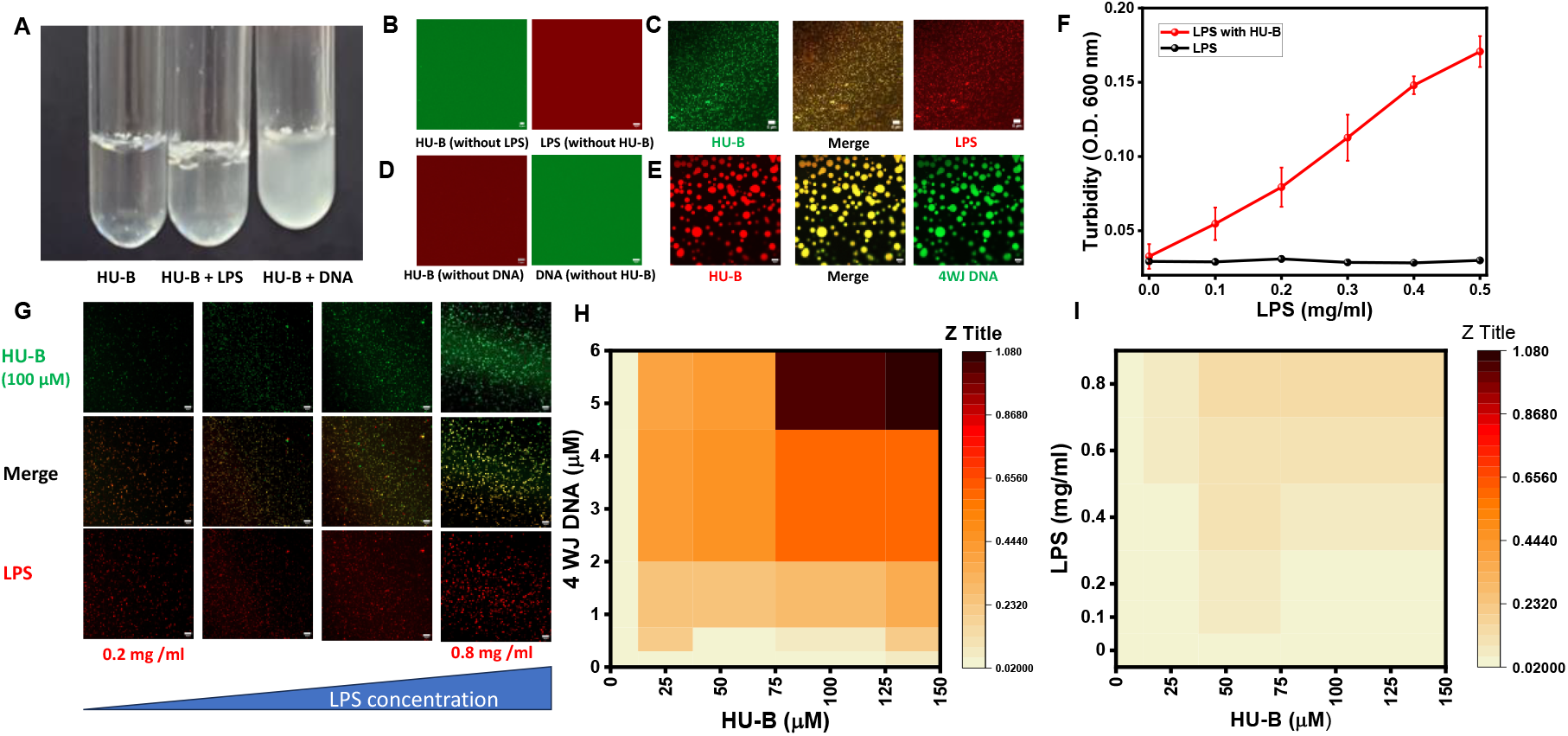
LPS and HU undergo biomolecular condensation. A) Images showing turbidity created in HU on addition of DNA or LPS. B, Control confocal microscopy images of HU (labelled with Alexa Fluor 488) and LPS (labelled with Alexa Fluor 594). C, Micrographs showing condensates created on mixing of HU-B (labelled with Alexa Fluor 488) and LPS (conjugated with Alexa Fluor 594). D, Control confocal microscopy images of HU (labelled with Alexa Fluor 594) and DNA (conjugated with Fluorescein). E, Micrographs showing condensates created on mixing of HU-B (labelled with Alexa Fluor 594) and DNA (conjugated with Fluorescein). F, Turbidity based examination of the effect of increasing LPS concentrations in presence of HU (black curve and in presence of HU-B (red curve). G, Micrographs of confocal microscopy showing the effect of increasing LPS (conjugated with Alexa Fluor 594). H, Phase diagrams of HU’s biomolecular condensation with (H) DNA and (I) LPS. All scale bars are of 5μM.

To compare the phase separation of HU-B with LPS and that of HU-B with DNA, we repeated the experiments shown in Figures 1B and 1C, using DNA. Figure 1D shows control confocal microscopic images of HU-B alone (100 μM), and DNA alone (6 μM), with no evidence of any phase separation. Figure 1E confirms our earlier reports regarding the occurrence of phase separation through mixing of HU-B and DNA, in a manner that leads to the formation of condensates that are both bigger and more well defined than the condensates formed by HU-B and LPS. We further verified the ability of LPS to modulate or increase the phase separation of HU-B, by increasing LPS concentrations and examining effects using both a turbidity-based assay (Figure 1F), and fluorescence microscopy (Figure 1G). Both experiments confirmed that higher concentrations of LPS elicit higher degrees of phase separation of HU-B, although condensates remain consistently smaller than condensates formed by equivalent concentrations of HU-B, and 4WJ-DNA. It may be noted, in passing, that HU exists in two isoforms in *E. coli*, HU-A and HU-B [4,22,23]. Throughout this article, we have used only HU-B, owing to its higher relevance to stress-related conditions, e.g., starvation [24], applicable to biofilms [25]; however, in Supplementary Figure S3 we also confirm the ability of the other HU isoform, HU-A, to phase separate with LPS and to form similar spherical biomolecular condensates that are rich in HU-A and LPS.

Next, we performed further comparisons of HU-LPS and HU-DNA condensates, using turbidity data as a proxy for the extent of phase separation, to constructs phase diagrams of increasing HU-B concentration as a function of either increasing DNA concentration (Figure 1H), or increasing LPS concentration (Figure 1I), using similar regimes and ranges of w/v concentration. In these phase diagrams, LPS concentrations are annotated in mg/ml, and 4WJ DNA concentrations are annotated in μM, since (as already mentioned) LPS has an average molecular weight of 75 KDa [17–19], causing 0.4 mg/ml LPS to be equivalent to approximately 5.33 μM LPS. On comparison of turbidities (represented as box color intensities in Figure 1H and 1I) for every combination of HU and DNA/LPS concentration, mole for mole, 4WJ DNA is clearly seen to cause phase separation of HU to a much greater extent than LPS. We hold that this is due to their being more HU-B binding sites on each copy of 4WJ DNA used [26], than on each copy of LPS, which contains only two sugar-phosphate moieties [8].

### Complex coacervation of LPS and DNA with HU

We next examined the effect of the presence of free LPS upon the biomolecular condensation of HU-B and DNA. In Figure 2A, LPS is demonstrated to be able to enter into condensates of HU-B and DNA to create complex coacervates that are rich in three molecular species; HU-B (blue), DNA (green) and LPS (red), all covalently labelled using different fluorophores. The calculation of colocalization coefficients shows that HU-B, LPS and DNA display significant colocalization in the same condensates, as shown in Supplementary Figure S4, implying that the three are able to together to form biphasic heterotypic condensates. In quality (if not in actual relative quantities), this mixture mimics the composition of EPS, since the EPS is rich in DNA and HU [27], and also contains LPS shed from the surfaces of *E. coli* cells [28]. In Figure 2B, we see the results of a turbidity-based experiment assessing the phase separation induced by a combination of LPS and DNA. The red curve plots turbidity for HU-B phase separating with 6 μM DNA. The light green curve plots turbidity for HU-B with 0.4 mg/ml LPS (∼5.33 μM). The blue curve plots turbidity for HU-B with both 6 μM DNA and 0.4 mg/ml (i.e., ∼5.33 μM) LPS. The curves clearly show the absence of any additive effect. Rather they show a negative effect, i.e., the presence of both LPS and DNA does not increase the extent of phase separation, but rather decreases the extent of phase separation. We believe that this is because LPS and DNA are already known to compete for the same sites in HU-B, and because LPS is poorer in its ability to phase separate with HU, than DNA.

**Figure 2.**
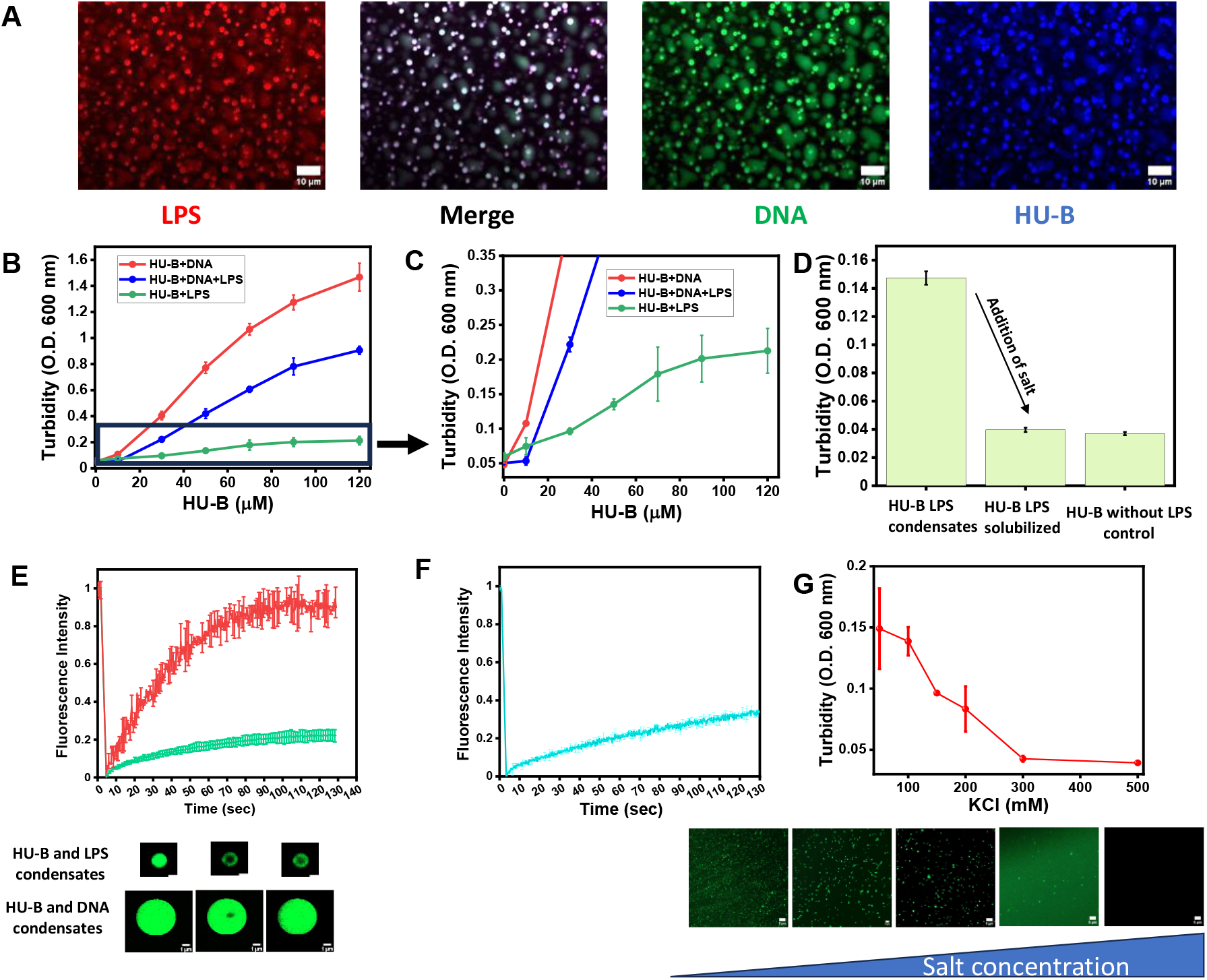
Effect of LPS on biomolecular condensation of HU and DNA, and the viscoelastic properties of condensates of HU with DNA or LPS. A, Confocal miscopy images of complex coacervation of HU (labelled with Alexa Fluor 647), LPS (labelled with Alexa Fluor 594) and (conjugated with Fluorescein). B, Turbidity based analysis of effect of complex coacervation of HU, LPS and DNA. C, expanded view of Figure 2B. D, Turbidity based assay showing reversibility of HU-LPS condensates on addition of salt. E, Averaged kinetics of fluorescence recovery after photo-bleaching (FRAP) of Alexa 488–labeled HU-B in condensates of HU-B and DNA (Red curve), and HU-B and LPS (green curve) with representative microscopy images of both experiments. F, Averaged kinetics of fluorescence recovery after photo-bleaching (FRAP) of Alexa 488–labeled HU-B, in condensates of HU-B and DNA spiked with LPS (final concentration 0.2 mg/ml). G, Turbidity based analysis of dependence on varying salt on HU-LPS biomolecular condensation. Microscopy images of the similar experiments are appended below. Fig. 2A has 10 μM scale bars, 2E has 1 μM scale bars and 2F has 5 μM scale bars.

We also verified that the observed HU-LPS condensates are not ‘HU-LPS aggregates’ by examining the reversibility of formation of the condensates through the addition of salt, which would be expected to reduce the binding of HU to LPS. Figure 2D shows that addition of 500 mM KCl to HU-LPS condensates formed in the presence of 150 mM KCl, reverses the turbidity obtained initially. Supplementary Figure S5 shows that condensates are no longer visible after the addition of 500 mM KCl.

### LPS can make HU condensates more gel like

There is another difference in the properties of HU-DNA and HU-LPS condensates, besides the differences in their average sizes noted above. Unlike HU-DNA condensates that rapidly wet surfaces, and also readily undergo fusion, HU-LPS condensates show no discernible wetting or fusion behavior (data not shown). Also, unlike HU-DNA condensates, which display rapid FRAP, i.e., recovery of fluorescence after photobleaching of flurophores attached to the components of the condensates (∼90% recovery in less than 60 seconds), HU-LPS condensates show only nominal recovery of fluorescence after photobleaching. This suggests that HU-LPS condensates are more gel-like in nature and less liquid-like than HU-DNA condensates (see the green curve in Figure 2E). Here, it must be noted that bleaching experiments with HU-LPS condensates that are typically very small posed significant experimental challenges. This caused us to resort to use of 10 % PEG 6000 and 0.8 mg/ml LPS, specifically for these FRAP experiments, in order to try and make HU-LPS condensates somewhat bigger and more stable over time, without appearing to lose water to the bulk solvent and ‘dry up’ rapidly. These attempts were made so that HU-LPS condensates could be subjected to photobleaching, with adequate scope remaining for the recovery of fluorescence after photobleaching, through diffusion of molecules from regions not covered by the laser, and not limited by the resolution of the experimental limits for specification of the region of interest targeted for bleaching.

Supplementary Figure S6 shows, as a control experiment, that no morphological changes occur in condensates of HU-B and LPS with the use of high concentrations of PEG 6000, while the red curve in Figure 2E shows FRAP data for condensates of HU-B and DNA under similar (control) PEG concentrations. This figure is shown while noting, in passing, that such conditions were not used in our previous paper reporting the discovery of the formation of LLPS condensates by HU and DNA [15]. The control experiments establish that the high PEG concentrations used do not cause the observed lower liquidity, since condensates of HU and DNA continue to show rapid FRAP at the very same high PEG concentrations used to obtain larger condensates of HU and LPS (10 % PEG), in which almost no FRAP is observed.

Since the high PEG concentrations used in the control FRAP experiments do not necessarily simulate macromolecular crowding in natural biofilms, we next decided to examine whether the addition of moderate levels of LPS to HU-DNA condensates leads to reduction of the liquidity of those condensates. Figure 2E shows that such ‘spiking’ of condensates of HU-B and DNA by free LPS (which simulates the presence of LPS shed by bacteria, within biofilms of HU and DNA) causes a significant reduction in liquidity, as measured through FRAP. This suggests that regions of biofilms that have high concentrations of cell-free LPS, as well as the immediate environments of LPS-bearing bacteria that are bound to either free, or DNA-bound, HU in biofilms, could be associated with less liquid-like and more gel-like characteristics. It may be noted that the viscoelastic nature of biofilms has already become a subject of interest amongst some investigators, and that modelling of the non-uniform viscoelastic nature of the extracellular matrix, over different regions of biofilms, has already been attempted [29,30].

### As with HU-DNA, HU-LPS condensates are destructible through the destruction of electrostatic interactions by salt

We have shown in the above sections that condensates of HU-B and LPS differ from condensates of HU-B and DNA in certain respects, despite the ability of the three species to form complex mixed coacervates. Here, we show that the molecular interactions driving the formation of the two types of condensates would appear to be similar. Just as HU-B’s interactions with DNA are governed by electrostatic forces, such forces also determine the formation of condensates of HU-B and LPS [15]. Figure 2F shows that increase in the concentration of KCl leads to reduction in turbidity, through reduction of the formation of condensates of HU-B and LPS. The same conclusion was also derived from confocal fluorescence microscopic experiments shown in the same figure.

### LPS-bearing *E. coli* cells adhere to (and enter) condensates of HU-B and DNA

In the above sections, we have demonstrated the compatibility of HU-B, DNA and LPS, in terms of their ability to form complex coacervates. Since HU-B binds to LPS through the same surfaces used by it to bind to DNA, as already shown earlier [8], it stands to reason that LPS-bearing *E*.*coli* cells may be anticipated to bind to ‘DNA-unoccupied’ sites on multimeric phase seperated forms of HU. We first initiated LLPS by addition of HU-B and DNA and added live *E. coli* cells expressing eGFP, on top of the sample, on the sample stage of the inverted microscope. In Figure 3 we see that, over time, cells appear to be titrated towards condensates of HU-B and DNA. In Figure 4A, maximum intensity projections of an experiment similar to the one shown in Figure 3 are shown. In Figure 3 and Figure 4A we can see that live GFP-expressing *E. coli* cells in stationary-phase adhere to (and enter) condensates of HU and DNA. We propose that the observed adherence, and entry, are the result of the titration of bacterial cells onto HU-B present on the surfaces of these condensates. We have already earlier shown that HU-B binds to the surfaces of *E. coli* cells and is able to cause the clumping of *E. coli* cells in both its free and DNA-bound forms [8].

**Figure 3.**
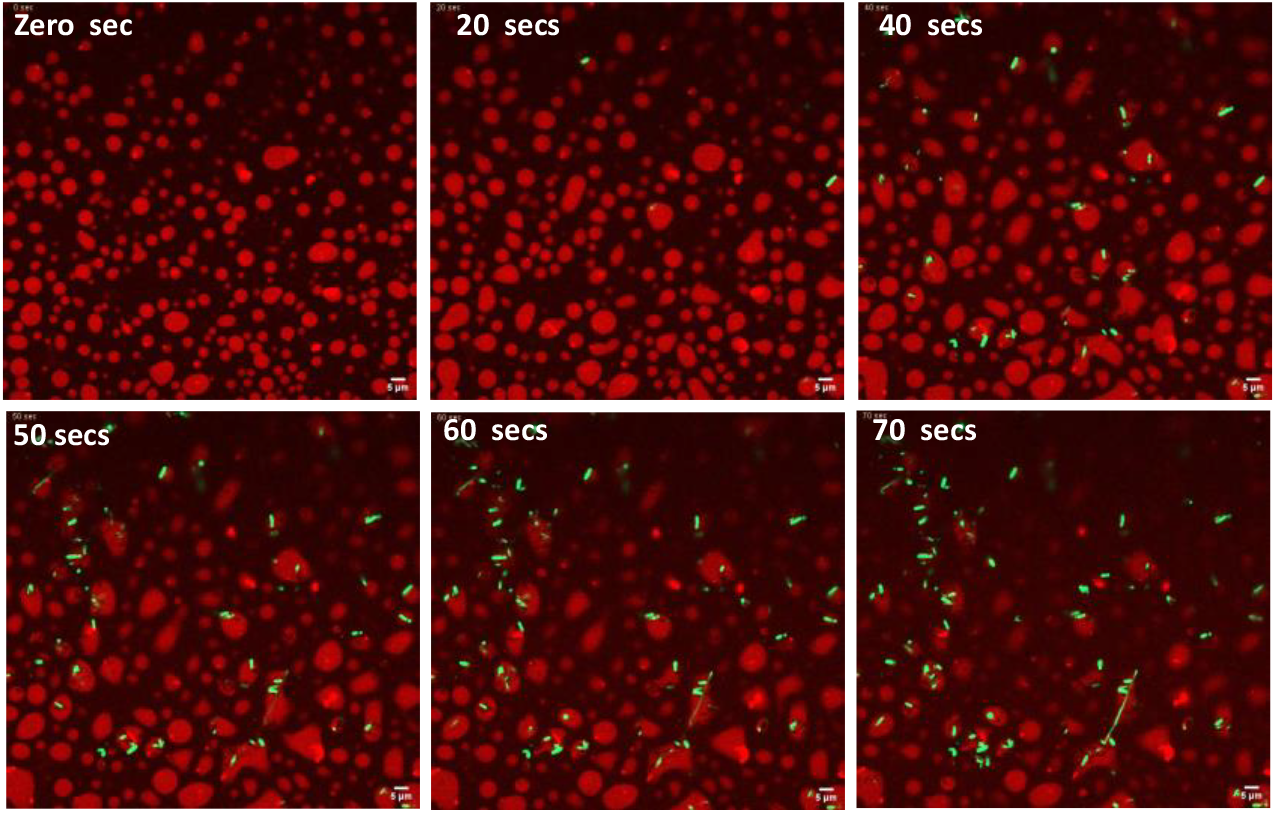
*E. coli* cells adhere to HU-DNA condensates. A time series of confocal microscopy showing live *E. coli* cells preferring to adhere to HU-DNA condensates.

**Figure 4.**
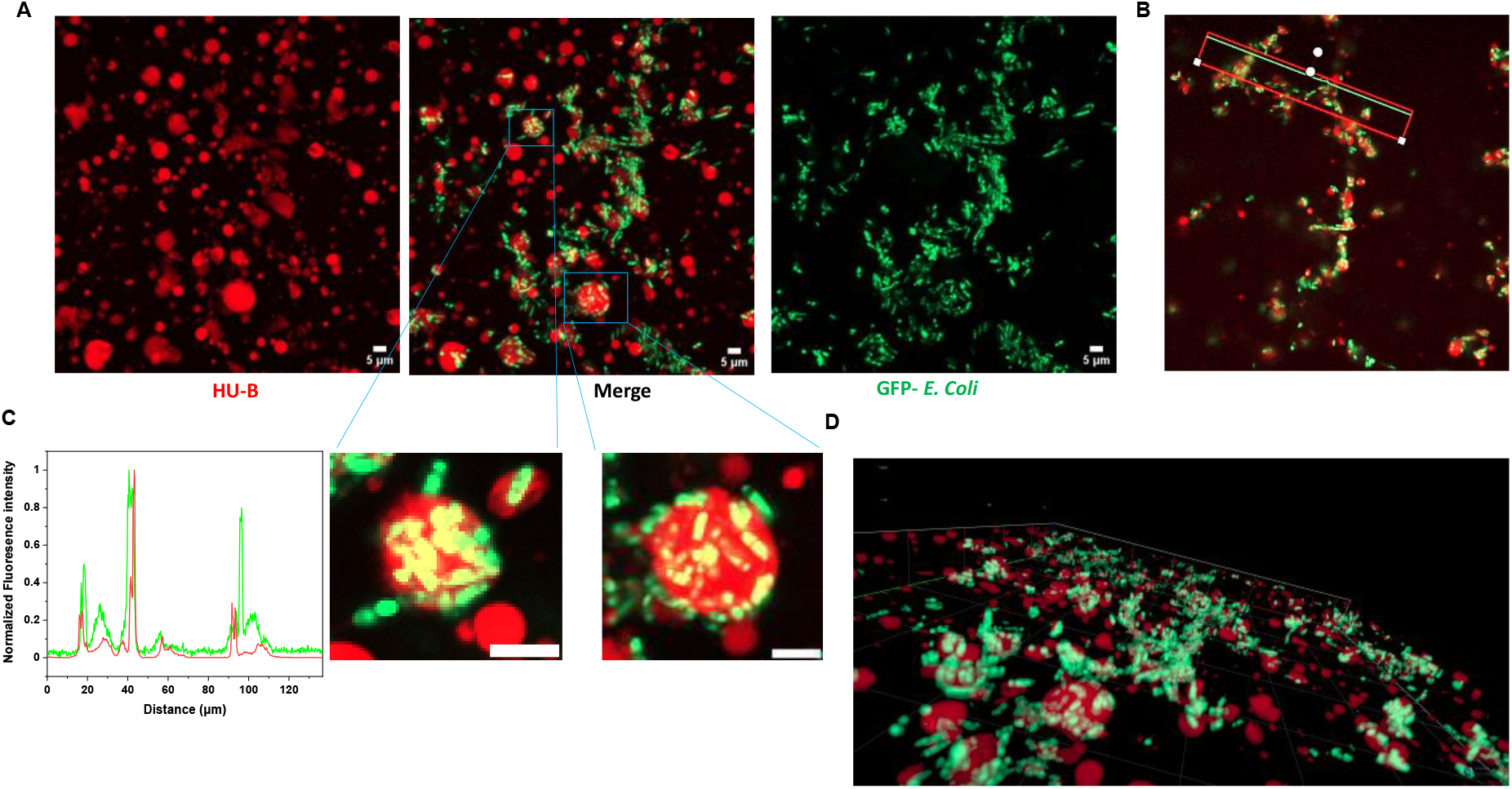
Arrangement of *E. coli* cells adhered to HU-DNA condensates. A, Maximum intensity projections of Z-stacks from confocal miscopy of *E. coli* cells adhering to HU-DNA condensates. HU-B is labelled via Alexa Fluor 647 and *E. coli* cells are fluorescing green because of GFP expression. B, Representation of a line that was drawn to calculate co-localization of condensates and *E. coli* cells. C, Line profile showing intensities of HU condensates (red) and *E. coli* (green), using the line shown in Figure 3B. D, A zoomed 3-D image showing cells coating the HU-DNA condensates. All scale bars are of 5 μM.

Figure 4A thus presents results with a simulacrum of a biofilm, in which bacteria with negatively-charged surfaces are able to become embedded within a matrix of DNA which is also negatively-charged, through the intercession of HU-B molecules that are able to act as a positively-charged glue. The variation of fluorescence intensity along a line drawn through the bacteria-associated condensates (shown in Figure 4B) is shown in Figure 4C, to demonstrate the colocalization of *E. coli* cells with condensates of HU-B and DNA, in that the presence of HU-B and DNA can be seen in the vicinity of most *E*.*coli* cells (although, of course, the presence of *E. coli* cannot be seen in the vicinity of all condensates formed by HU-B with DNA. In Figure 4C, the green and red intensities of *E. coli* and HU-B, respectively, either overlap (for cells buried within condensates) or adjoin each other (for cells adhering to condensates). Figure 4D shows a zoomed-in section of a 3-D representation created through the overlapping of a stack of images collected along the z-axis, in a series of confocal microscopic experiments. Supplementary video 1 shows a video created out of merged channels of Z stacks from the experiment shown in Figure 3, with every stack appearing as a frame in the video. This figure and the video confirm that bacterial cells adhere to (and enter) condensates of HU and DNA. In addition, Suppmentary video 2 also shows frames of a single condensate as a function of viewing depth. In this video, it can be seen very clearly that some *E. coli* cells are attached to the condensate while many *E. coli* cells are located in the interiors of the condensate.

## Discussion

Biofilms are a way of life for microbes in the wild, and also inside the bodies of other organisms. Inside organisms, biofilms that form on the surfaces of vessels carrying blood, or food, can potentially resist the entry of antibiotics and thus become hosts for populations of bacteria that can display proliferation after a course of antibiotics has concluded, and also act as nurseries for the generation of antibiotic-resistant populations of mutant bacteria. Recent findings, such as those by Seviour *et al*., highlight the significance of surface proteins like Brosi-A1236 in *Candidatus Brocadia* in phase separation and cell-cell adhesion for biofilm formation [31]. Staphylococcal Bap protein has also been shown to be able to forms amyloids (after maturation of LLPS condensates) and act as an extracellular matrix scaffold in biofilms [32]. Such findings prompted us to explore the role of LLPS in *E. coli* biofilm formation involving the NAP, HU.

In this article we demonstrate for the first time that liquid-liquid phase-separated condensates of a DNA binding protein can act as a simulacrum of biofilms. We have used cellular concentrations of HU [33]. DNA concentrations in the EPS of biofilms are known to fluctuate significantly, ranging from 1 mg/ml to 40 mg/ml [34]. To the best of our knowledge LPS concentrations have never been determined experimentally, therefore we have used a nominal concentration of 0.2 mg per ml in most experiments. This being said, it must be appreciated that the EPS inside a biofilm is highly dynamic, and that metabolites inside biofilms are known to exist in different gradients and can be present at concentrations many folds higher at one point in the biofilm, than at another. [35] Using such proxy concentrations, we show that HU, one of the most abundant and crucial protein components of the extracellular matrix in any *E. coli* biofilm, undergoes complex coacervation with DNA and/or LPS to create condensates whose liquidity is determined by LPS content, with LPS acting like a mimic of DNA and binding to DNA-binding sites on HU through electrostatic interactions. It is known that *E. coli* cells shed LPS into the extracellular matrix. We show, using fluorescence recovery data after photobleaching, that LPS makes phase separated condensates of HU-B and DNA more gel-like. Presumably, this allows cells to stick to surfaces with high fluid dynamics and turbulence, such as wet rocks, gut lining and the inner surfaces of blood vessels. Taking together our previous articles and this article, we show that HU probably plays a central role in phase separating with DNA both within bacterial cells (to form a phase-separated genomic nucleoid), and outside bacterial cells (to form the body of a biofilm). The phase separation of the proteinaceous and extracellular polymeric substances in a biofilm provides benefits such as the exclusion of nucleases, or proteases, and other molecules (such as antibiotics) that could affect cell viability, by slowing down diffusion in a manner modulated by LPS concentration.

LPS is known to be capable of existing in a phase-separated state in the outer membrane of *E. coli* and to create LPS raft-like structures that are most likely involved in immunoregulation [36]. There exists a possibility that phase separated LPS on bacterial cell surfaces facilitates the interaction of HU-DNA-LPS in the EPS and allows adhesion of cells to each other and to the EPS. In the end, we show that *E. coli* cells move into condensates of HU-B and DNA, presumably because HU-B binds to cell-displayed LPS and because condensates contain a higher concentration of HU-B than the bulk solvent, thus allowing HU-DNA condensates to essentially act like kernels for initiating biofilm formation. In totality, in this article, we have shown that biofilms benefit from the phase separation of DNA with DNA-binding proteins. We have also shown that addition of LPS to phase-separated structures can make the EPS more gel like and aid in cellular embedment.

## Data availability

All the data for this manuscript is contained in the article above, and the supplementary information file.

## Acknowledgments

We thank the Government of India for financial support (for equipment/consumables) extended through IISER Mohali. Gurmeet Kaur, Arpita sarkar and Soumyajit Kar are acknowledged for assistance rendered during their training on the imaging systems.

## Funding and additional information

We thank the Ministry of Education, Government of India for a Centre of Excellence grant in Protein Science, Design and Engineering (MHRD-14-0064), awarded to P.G., the TATA Transformation Prize Grant, awarded to P.G., & the DST-FIST grant awarded to the Department of Biological Sciences, IISER Mohali. A. G. thanks DBT (India) for a research fellowship.

## Experimental procedures

### 4WJ DNA

4WJ DNA was generated *in vitro* by annealing of four different oligos (purchased from Merck) in equimolar amounts to create a Holliday junction–like (cruciform) structure. The nucleotide sequences of four oligos used were as follows:

5’ - CCCTATAACCCCTGCATTGAATTCCAGTCTGATAA3’,

5’-GTAGTCGTGATAGGTGCAGGGGTTATAGGG-3’,

5’-AACAGTAGCTCTTATTCGAGCTCGCGCCCTATCACGACTA-3’,

5’-TTTATCAGACTGGAATTCAAGCGCGAGCTCGAATAAGAGCTACTGT-3’.

4WJ DNA was used at a concentration of 3 μM, unless stated otherwise.

### Turbidity assays

All components were mixed on the bench and then immediately transferred to a 96-well plate for absorption measurements in a BMG Labtech POLARstar Omega plate reader. Absorption readings were taken at 600 nm at 37 °C.

### Reversibility of HU and LPS condensates

Phase separation was induced, as usual, by adding LPS to HU. Formation of condensates was verified by measuring turbidity and by visualization using confocal fluorescence microscopy. To these preformed condensates, KCl was added from a stock of 3M, the dissolution of condensates was verified by the same assays as described above.

### Microscopy

Confocal fluorescence microscopic imaging was performed using a ZEISS LSM 980 Elyra 7 super-resolution Microscope using a ×63 oil-immersion objective with a numerical aperture of 1.4, coupled with a monochrome cooled high-resolution AxioCamMRm Rev. 3 FireWire(D) camera. Samples for microscopy were prepared by mixing on the bench and were imaged immediately thereafter, using makeshift chamber slides. For samples with proteins, fluorescently labelled proteins were spiked into the unlabeled proteins; spiking was done with 1 %; 99 % of the protein was accounted for by unlabeled protein with no cysteine mutation. All images were analyzed using Fiji ImageJ.

### Protein purification and concentration estimation

Protein expression and concentration protocols have been detailed in previous articles [8,15]. Briefly, pQE-30 plasmids (Qiagen) containing the gene encoding HU-B F47W were transformed into XL1-Blue cells, and expressed protein was purified using Ni-NTA chromatography, followed by cation exchange chromatography. The protein used in this article was a point mutant (HU-B F47W) of the wild type HU-B protein. HU-BF7W was used because wildtype HU has no tryptophan in it, for reasons elaborated earlier [37]. Without tryptophan in its sequence, it is difficult to calculate protein concentrations accurately therefore, we have used the mutant which we have shown to not cause effect on DNA binding or phase separation ability of HU [8,15].

### FRAP assay

FRAP assays were carried out using a Zeiss LSM 980 microscope. In all cases HU-B was bleached using a 488 nm laser line. For bleaching, 100% intensity of laser was used and 100 iterations were made in a circle of 0.5 μm diameter. Experiments were done in triplicates and fluorescence intensities were plotted after normalization.

### Fluorescent labelling

LPS with covalently conjugated Alexa Flour 594 was purchased from Merck (cat. No. L23353). A cysteine mutant of HU, described previously [15], was used to label HU with Alexa fluor 647 C_5_ Maleimide or Alexa Fluor 488 C_5_ Maleimide. The procedure for labelling involved overnight incubation of 100 μM HU-B with 300 μM of TCEP (Tris(2-carboxyethyl) phosphine hydrochloride, and 200 μM of dye. Ultracentrifugation was done to remove free dye.

### Imaging of cells adhering to condensates of HU and DNA

Condensates of HU-B and DNA were made *in vitro* with spiking of either Alexa 594-labelled HU or Alexa Fluor 647 labelled HU. The condensates were then visualized using LSM 980 confocal microscope. On top of the coverslip, containing HU-B and DNA condensates, stationary phase *E. coli* cells were added. An aliquot of the mixture was visualized immediately 3-D rendition of the z stacks was prepared using Zen blue edition software. The line profile for fluorescence intensities was also prepared using Zen blue edition software, and extracted raw values were plotted using origin pro 2020b.

### Colocalization image analysis

Images were loaded into ImageJ, followed by the application of max entropy thresholding. The Pearson’s correlation coefficient and Mander’s overlap coefficients (MI and M2) were then computed using the JaCOP plugin.

### Statistical analysis

All experiments were done in triplicates. The data for mean values was plotted along with standard deviation. The statistical significance analysis was performed using one-way ANOVA analysis of variance test, using the software Origin Pro 2021b. All data analysis, data fitting, and data plotting were performed using Origin Pro 2021b.

